# Non-invasive diagnostic method to objectively measure olfaction and diagnose smell disorders by molecularly targeted fluorescent imaging agent

**DOI:** 10.1101/2021.10.07.463532

**Authors:** Dauren Adilbay, Junior Gonzales, Marianna Zazhytska, Paula Demetrio de Souza Franca, Sheryl Roberts, Tara Viray, Raik Artschwager, Snehal Patel, Albana Kodra, Jonathan B. Overdevest, Chun Yuen Chow, Glenn F. King, Sanjay K. Jain, Alvaro A. Ordonez, Laurence S. Carroll, Thomas Reiner, Nagavarakishore Pillarsetty

**Affiliations:** Department of Radiology, Memorial Sloan Kettering Cancer Center, New York, NY, USA; Department of Surgery, Memorial Sloan Kettering Cancer Center, New York, NY, USA; Mortimer B. Zuckerman Mind, Brain and Behavior Institute, Columbia University, New York, NY 10027, USA; Department of Genetics and Development, Columbia University Irving Medical Center, Vagelos College of Physicians and Surgeons, Columbia University, New York, NY, 10032, USA; Department of Otolaryngology- Head and Neck Surgery, Columbia University Irving Medical Center, Vagelos College of Physicians and Surgeons, Columbia University, New York, NY, 10032, USA; Institute for Molecular Bioscience, The University of Queensland, St Lucia, QLD 4072, Australia; Australian Research Council Centre of Excellence for Innovations in Peptide and Protein Science, The University of Queensland, St Lucia, QLD 4072, Australia; Center for Infection and Inflammation Imaging Research, Johns Hopkins University School of Medicine, Baltimore, MD, USA; Department of Pediatrics, Johns Hopkins University School of Medicine, Baltimore, MD, USA; Russell H. Morgan Department of Radiology and Radiological Sciences, Johns Hopkins University School of Medicine, Baltimore, MD, USA; Department of Radiology, Weill Cornell Medical College, New York, NY, USA; Chemical Biology Program, Memorial Sloan Kettering Cancer Center, New York, NY, USA

## Abstract

The sense of smell (olfaction) is one of the most important senses for animals including humans. Despite significant advances in the understanding mechanism of olfaction, currently, there are no objective non-invasive methods that can identify loss of smell. Covid-19-related loss of smell has highlighted the need to develop methods that can identify loss of olfaction. Voltage-gated sodium channel 1.7 (Na_V_1.7) plays a critical role in olfaction by aiding the signal propagation to the olfactory bulb. We have identified several conditions such as chronic inflammation and viral infections such as Covid-19 that lead to loss of smell correlate with downregulation of Na_V_1.7 expression at transcript and protein levels in the olfactory epithelium. Leveraging this knowledge, we have developed a novel fluorescent probe Tsp1a-IR800 that targets Na_V_1.7. Using fluorescence imaging we can objectively measure the loss of sense of smell in live animals non-invasively. We also demonstrate that our non-invasive method is semiquantitative because the loss of fluorescence intensity correlates with the level of smell loss. Our results indicate, that our probe Tsp1a-IR800, can objectively diagnose anosmia in animal and human subjects using infrared fluorescence. We believe this method to non-invasively diagnose loss of smell objectively is a significant advancement in relation to current methods that rely on highly subjective behavioral studies and can aid in studying olfaction loss and the development of therapeutic interventions.

## Introduction

The sense of smell (Olfaction) is critical for survival of animals including humans because smells convey information about the food, environment and serves as tools for inter- and intra-species communication. The loss of sense of smell (Anosmia), resulting from Covid-19 associated neurological sequelae has captured several headlines over past two years and has highlighted the need for gaining deeper mechanistinc understanding and developing tools to probe such disorders [1, 2]. This also has shed light on general olfaction disorders associated with several diseases because anosmia has a tremendous negative impact on the individual quality of life and public health [3, 4]. A staggering 13.3 million adults in US were found to have smell disorders, including 3.4 million with severe hyposmia/anosmia. These studies has been conducted before the COVID-19 pandemic, and these numbers can change drastically with millions of infections and due to long Covid-19 [5-9]. Despite such a fundamental importance in biology and high prevelance of Anosmia, we do not have an objective method to measure perception of smell. The perception of smell like other senses is subject reported and therefore a non-independent measure. This also implies, we do not have any objective tools to detect presence or absence of olfaction in animal models. This has resulted in reliance on behavioral studies that are not only challenging but are indirect and highly subjective. This has impaired investigations in animal models, and, thereby, developing new therapeutic interventions for anosmia. Therefore, there is an urgent and compelling need to develop tools that can measure sense of smell in human and animal models.

To develop a diagnostic method for olfaction, we explored the potential of voltage gated sodium channel 1.7 (Na_V_1.7). We reasoned that sodium channels (Na_V_1.1-1.9) are crucial members in neurotransmission and aid in propagation of action poetential/impulse along the nerve bundles and dysfuntion of these channels might lead to impaired olfaction. Na_V_1.7 encoded by SCN9A gene is expressed predominantly in peripheral sensory neurons and the axons of human olfactory sensory neurons (OSNs) and plays a critical role in olfaction among mammals [10]. In OSNs, Na_V_1.7 is critical for transmitting olfactory cues provided by olfactory receptor activation to higher-order neurons in the brain. It has been established that dysfunction of Na_V_1.7 resulting from mutations in the SCN9A gene encoding Na_V_1.7 or loss of these sodium channels at the level of olfactory bulb and olfactory epithelium (OE) results in smell disorders [10]. The presence of Na_V_1.7 in superficial layer of the OE presents an unprecedented opportunity to develop optical probes that can non-invasively image expression of Na_V_1.7. By objectively measuring changes in Na_V_1.7 expression in OE we can potentially identify smell disorders in patients and animals by measuring changes.

In the current manuscript we first establish that Na_V_1.7 is a bona fide target for olfaction by demonstrating that conditions such as viral infections or inflammation resulting from allergies or drugs, that leads to the loss of odorant perception, results in diminished expression of Na_V_1.7 in the superficial layer of the OE. We also demonstrate that SARS-CoV-2-infected mice, hamsters and humans have diminished Na_V_1.7 expression in the OSN using either immunohistochemistry (IHC), single cell and/or bulk RNA-seq. Taking advantage of a potent and subtype-selective peptide Tsp1a isolated from Peruvian tarantula that targets Na_V_1.7, we have developed fluorescent imaging agents that can aid visualization of peripheral nerves *in vivo* without any side-effects [11, 12]. Using a fluorophore (IR800) that is better suited for *in vivo* imaging applications and clinical translation, we have genearated and characterized Tsp1a-IR800 that is specific for Na_V_1.7. We demonstrate that Tsp1a-IR800 can be used to non-invasively image Na_V_1.7 on OE in mice models using epifluorescence imaging and that methimazole induced chronic inflammation anosmia can be non-invasively detected by our imaging probe Tsp1a-IR800. In non-human primate (NHP) imaging studies with Tsp1a-IR800 (*i*.*v*.) the fluorescent intensity of the NHP olfactory epithelium was significantly higher compared to surrounding tissues. Our IHC studies indicate human OE expresses Na_V_1.7 and its expression is downregulated in Covid-19 patients. Taken together, our findings demonstrate that our fluorescent imaging probe Tsp1a-IR800 targeting Na_V_1.7 channels can serve as a non-invasive imaging tool for anosmia.

## Materials and methods

### Synthesis of Tsp1a-IR800

Synthetic Tsp1a was prepared as in our previous report [11, 13]. Tsp1a-IR800 was synthesized using Tsp1a peptide (0.71 mM, 250 μg in 100 μL of H_2_O), which was diluted in a 25-mM Tris-buffered aqueous solution (Fig. 2). To this reaction mixture, 20 μL of a 50-mM solution of L-ascorbic acid in H_2_O and 20 μL of a 50-mM aqueous CuSO_4_ solution in H_2_O, were added. This step was followed by the addition of 50 μg (44 nmol) of IR800 azide in 50 μL of acetonitrile, after which the reaction mixture was stirred at room temperature in the dark for 4 h. The crude mixture was separated via reversed-phase (RP) HPLC, yielding 44 μg (10 nmol; 26%) of Tsp1a-IR800. The final analytical RP-HPLC purification showed 96% purity. LCMS-ES (ESI+), m/z calculated for Tsp1a_Pra_IR800, [C_207_H_293_N_48_O_57_S_10_] 4682.87, [C_207_H_293_N_48_O_57_S_10_+3H]^3+^ 1563.09, found [M+3H]^3+^ 1564.10, [C_207_H_293_N_48_O_57_S_10_+4H]^4+^ 1172.00, found [M+4H]^4+^ 1173.20, [C_207_H_293_N_48_O_57_S_10_+5H]^5+^ 938.16, found [M+5H]^5+^ 939.00.

### Animal works

All animal care and procedures were approved by the Animal Care and Use Committees (IACUC) of Memorial Sloan-Kettering Cancer Center.

#### Mice

Hsd:athymic female mice Nude-Foxn1^nu^ (6–8 weeks old) and *Chlorocebus aethiops* primates (5-8 months) were used in the study. We first assessed the possibility of visualizing the olfactory nerve in normosmic mice using Tsp1a-IR800. Nine athymic nude mice were divided into three groups. The main group was intravenously injected with Tsp1a-IR800 (1 nmol, 100 μL PBS), the control group was injected with phosphate-buffered saline (PBS), and a third group was injected with a combination of Tsp1a-IR800 and unmodified Tsp1a (blocking agent).

#### Mice infected with SARS-CoV-2

Heterozygous K18-hACE2 C57BL/6J 8-week old male mice (strain: 2B6.Cg-Tg(K18-ACE2)2Prlmn/J, Jackson Laboratory), received 8.4 × 105 TCID50 of SARS-CoV-2/USA/WA1/2020 intranasally, after induction of anesthesia with ketamine hydrochloride and xylazine (PMID: 35304580). Six days after infection, the animals were sacrificed with isofluorane overdose. The tissues were harvested and fixed in 10% formalin for 48 hours.

#### Hamsters

LVG Golden Syrian hamsters (Mesocricetus auratus) were treated and euthanized in compliance with the rules and regulations of IACUC under protocol number PROTO202000113-20-0743. Only male hamsters were used for experiments. All experiments were performed on dissociated cells prepared from whole olfactory epithelium tissue.

#### Non-human primates

The non-human primates (2 males, 1 female) were intravenously injected with Tsp1a-IR800 (200 μg/kg) and sacrificed after 120 minutes. Tissues of the olfactory epithelium, bulb, muscle, and brain were isolated and resected during necropsy.

#### Olfactory ablation

Methimazole treatment was performed to ablate the olfactory nerve as described elsewhere [14]. Mice were injected with methimazole dissolved in isotonic saline (50 mg/kg body weight) via intraperitoneal administration, on days 0 and 3. *In vivo* fluorescence imaging was performed on day 8 after the first injection. We have used 3 groups of mice, treated with methimazole, normosmic mice, and a blocking group. After imaging, all mice were sacrificed, and organs of interests were dissected.

#### Behavioral experiment

The buried food test was performed as described [15]. Briefly, mice were fasted overnight by removing all chow pellets from the cage 18–24 h before the test. On the test day, a mouse was transferred to the clean cage containing 3 cm bedding, with the hidden cookie in one of the random corners 2 cm deep. The observer retreats to the observation station and starts the timer and stops the stopwatch when the subject mouse finds the buried food (Fig. 4A). A subject was considered to have uncovered the cookie when it starts to eat, usually holding the food with forepaws.

#### Fluorescence Imaging of mice

All mice were anesthetized using an intraperitoneal injection of a ketamine/xylazine cocktail and *in vivo* epifluorescence images obtained using an IVIS Spectrum imaging system (PerkinElmer) using filters set for 745 nm excitation and 810 nm emission. Animals were sacrificed and olfactory nerve/bulb, muscle, heart, spleen, kidney, and liver were removed and imaged *ex vivo* using the IVIS system. Autofluorescence was removed through spectral unmixing. Semiquantitative analysis of the Tsp1a-IR800 signal was conducted by measuring the average radiant efficiency (in units of [p/s/cm^2^/sr]/[μW/cm^2^]) in regions of interest (ROIs) that were placed on the ROEB.

#### Tabletop fluorescence microscopy

Olfactory epithelium tissues of mice intravenously injected with PBS (control), Tsp1a-IR800 or Tsp1a-IR800/Tsp1a formulation were stained with a solution 10 μg/mL Hoechst 33342 in PBS and placed over a coverslip glass slide (48 × 60 mm no. 1 thickness, Brain Research Laboratories, Newton, MA). Images were acquired using a laser scanning inverted-stand confocal microscope (Leica TCS SP8, Germany) using 725 nm laser excitation for IR800 (red) and 405 nm for Hoechst (blue).

#### Dissection of olfactory bulb and epithelium in mice

Mice were anesthetized with ketamine/xylazine (100 and 20 μg/g body weight, respectively), perfused transcardially with 0.1 M phosphate buffer (PB) followed by a PB buffered fixative containing 4% paraformaldehyde (PFA) as described by Lin et al [16]. The dissection is divided into two components, the removal of the brain from the body and the removal of the skin and lower jaw. Under magnification, we removed the hard palate and the bones covering the brain, keeping nasal bones in place. The head was fixed in 4% PFA overnight. Tissues were processed, decalcified and paraffin-embedded using a standard protocol.

#### Single cell RNA-seq

Cells were dissociated according to the Worthington Papain Dissociation System by incubating fresh olfactory tissue with papain for 40 min at 37 °C. Library preparation was performed accordingly to Chromium Single Cell 5′ v.2 Protocol and sequenced on NextSeq550. Cell Ranger pipelines were used to generate fastq files which subsequently were aligned against MesAur1.0/WuhCor1 genomes. Systemic biases and background noise were removed with Cellbender; resulted h5 matrixes were loaded and analyzed with Seurat. Cells with more than 400 UMIs, expressed 500 and 6000 genes and less than 5% of mitochondrial genes were kept for further analysis. Identified 13 clusters were visualized with UMAP and annotated using known marker genes for each cell type. Differential expression analysis was performed using the default two-sided non-parametric Wilcoxon rank sum test with Bonferroni correction using all genes in the dataset.

#### Bulk RNAseq from human olfactory epithelium

RNA was extracted using Direct-zol RNA kits from Zymo Research. 50ng-1ug of total RNA was used to prepare DNA libraries with Truseq RNA Library Prep Kit v2 followed by 75 HO paired-end and multiplexed sequencing. Reads were aligned to human genome (hg38), *Mesocricetus auratus* (MesAur1.0) and SARS-CoV-2 (wuhCor1) using subread and the raw read counts were assembled using feature counts pipeline.

#### Immunohistochemistry and quantification of Na_V_1.7 expression

Na_V_1.7 staining was performed at the Molecular Cytology Core Facility of MSK using a Discovery XT processor (Ventana Medical System, Tucson, AZ). We used an anti-Na_V_1.7 antibody [N68/6] (NeuroMab) that binds to both human and mouse Na_V_1.7 (0.5 μg/mL). Paraffin-embedded formalin-fixed 4 μm sections were deparaffinized with EZPrep buffer. For IHC, a 3,3′-diaminobenzidine (DAB) detection kit (Ventana Medical Systems, Tucson, AZ) was used according to the manufacturer’s instructions. Sections were counterstained with hematoxylin and eosin (H&E) and coverslip using Permount (Fisher Scientific, Pittsburgh, PA).

Na_V_1.7 quantification was performed on digitalized slides. The threshold for signal intensity in the DAB (brown) and H&E (blue, representing all tissue area) channels was determined via an automated script using ImageJ analysis software. Color deconvolution was used to separate blue and brown signals and the threshold values were kept constant: 0–114 for DAB and 0–235 for H&E. The relative Na_V_1.7 positive area was calculated by dividing the brown (DAB) area by the blue (total tissue area).

#### Fluorescence imaging of non-human primate tissues

After intravenous injection of Tsp1a-IR800, animals (*C. aethiops*) were euthanized and several tissues including olfactory epithelium, olfactory bulb, muscles and brain were harvested and dissected. For imaging using Quest Camera, tissues were placed on a special black paper to minimize reflection. The camera of the Quest NIR imaging system was placed above the tissues at 15 cm distance. The same settings were used for typical imaging procedures (30 ms exposure time, 100% excitation power, gain: 25.5 dB, at 24 frames per second). Analysis and quantifications were performed on Quest QIFS Researchtool (Quest Medical Imaging, Middenmeer, Netherlands). Regions of interests (ROI) were drawn on the dissected tissues using polygon instrument. Total 4 ROIs were selected to encompass the olfactory epithelium, muscle, olfactory bulb and brain of the non-human primate.

#### Statistical analysis

Statistical analysis of Na_V_1.7 expression was performed using R v3.6.3 (R Core Team - 2020, R Foundation for Statistical Computing, Vienna, Austria) and GraphPad Prism 9 (GraphPad Software, San Diego, CA). Student’s *t*-test was used to compare Na_V_1.7 expression (ratio Nav1.7 expression/Total tissue area) between different groups of mice. Pearson correlation coefficient was used to examine the correlation between the time in food buried test and radiant efficiency. Results with a p-value of 0.05 were considered statistically significant. Data points represent mean values, and error bars represent standard deviations.

## Results

### Na_V_1.7 is abundantly expressed in the region of the olfactory epithelium and bulb (ROEB) of normosmic mice

We isolated and dissected olfactory bulb and epithelium regions from normosmic nude mice, and used immunohistochemistry (H&E and anti-Na_V_1.7 antibody staining) to identify areas of high Na_V_1.7 expression. Consistent with previous reports [10, 17], we found that Na_V_1.7 is moderately expressed in layers of primary OSNs, and highly expressed in olfactory nerve bundles located in lamina propria, including all the paths towards the olfactory bulb (Fig. 1B). Olfactory bulbs abundantly express Na_V_1.7 on peripheral areas, which corresponds to the olfactory nerve layer (ONL). The deeper, glomerular layer located just after the ONL corresponds to terminal synapses of OSN axons with the dendrites of mitral and tufted cells, and was lightly stained for Na_V_1.7 (Fig. 1B).

**Figure 1.**
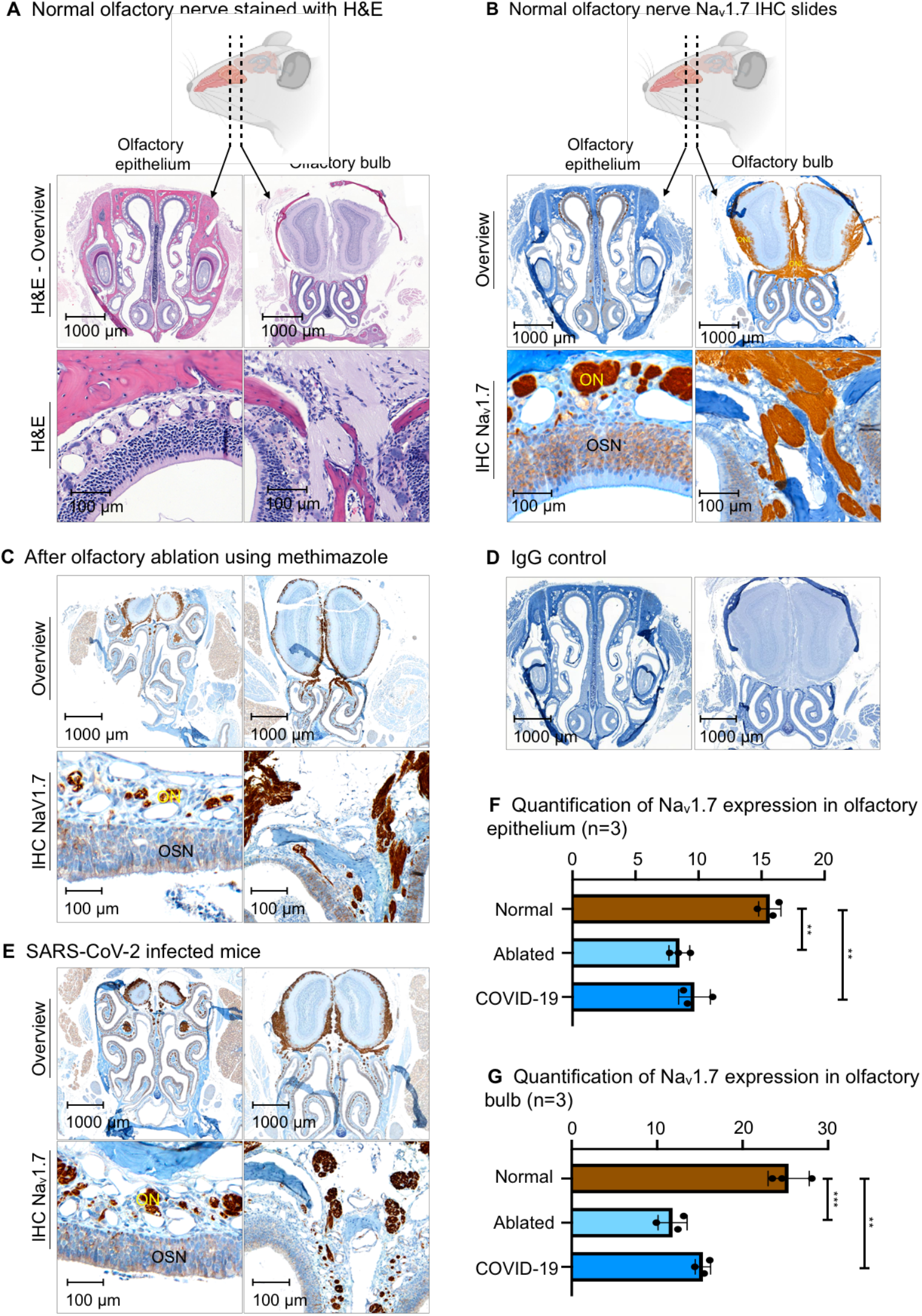
Histological slides of the olfactory bulb and olfactory epithelium of normosmic mice, mice with olfactory ablation, and a mouse infected with COVID-19. (A) H&E stain of normosmic mouse. (B) Na_V_1.7 IHC slide of normosmic mouse. (C) IHC slide of the mouse after olfactory ablation. (D) IHC slide of olfactory tissue with IgG isotype primary antibody. (E) IHC slides of a mouse with SARS-CoV-2 infection. (F) Quantification of Na_V_1.7 expression in the olfactory epithelium of 3 mice. (G) Quantification of Na_V_1.7 expression in olfactory bulb of 3 mice. (** P ≤0.01; *** P ≤ 0.001; ON – olfactory nerve bundles; OSN – olfactory sensory neurons; ONL – olfactory nerve layer)

#### Na_V_1.7 expression is reduced in chemically-induced anosmia and COVID-19 disease models

The expression of Na_V_1.7 was significantly diminished after olfactory ablation using methimazole and also in SARS-CoV-2 infected mice (Fig. 1C, E). Of the total tissue area in the olfactory epithelium of normosmic mice, 15.7% was the Na_V_1.7-positive area, compared to 8.5% in olfactory-ablated mice and 9.7% in SARS-CoV-2 infected mice (Fig. 1F). The ONL of the olfactory bulb also showed a decrease in Na_V_1.7 expression in both olfactory-ablated and SARS-CoV-2 infected mice (Fig. 1G).

### RNA sequencing data reveals temporal downregulation of SCN9A gene expression in OSN cells from hamsters infected with SARS-CoV-2 and correlates with loss- and gain of olfactory function

We performed bulk and single-cell RNA-seq of SARS-CoV-2 infected and mock hamsters’ olfactory epithelium tissues at 1-, 3- and 10-days post-infection (dpi). For scRNA-seq we analyzed 68.951 cells and identified 13 cell subtypes using previously described markers [18]. SCN9A was predominantly expressed in OSN’s and olfactory glia with minimal to zero expression in other cell subtypes in the OE. In SARS-CoV-2 infected hamsters we observed a 3-fold drop in the expression of the SCN9A gene transcripts in OSNs at 3 dpi, and full restoration of expression at 10 dpi with bulk RNA-seq analysis (p<0.001). These changes in SCN9A transcripts can be followed on UMAP clustering maps that show temporal downregulation of SCN9A. At 3 dpi, the transcript levels are significantly diminished compared to control as well as 1 dpi and are restored to pre-infection levels 10 dpi (Figure 2a, b, c).

**Figure 2.**
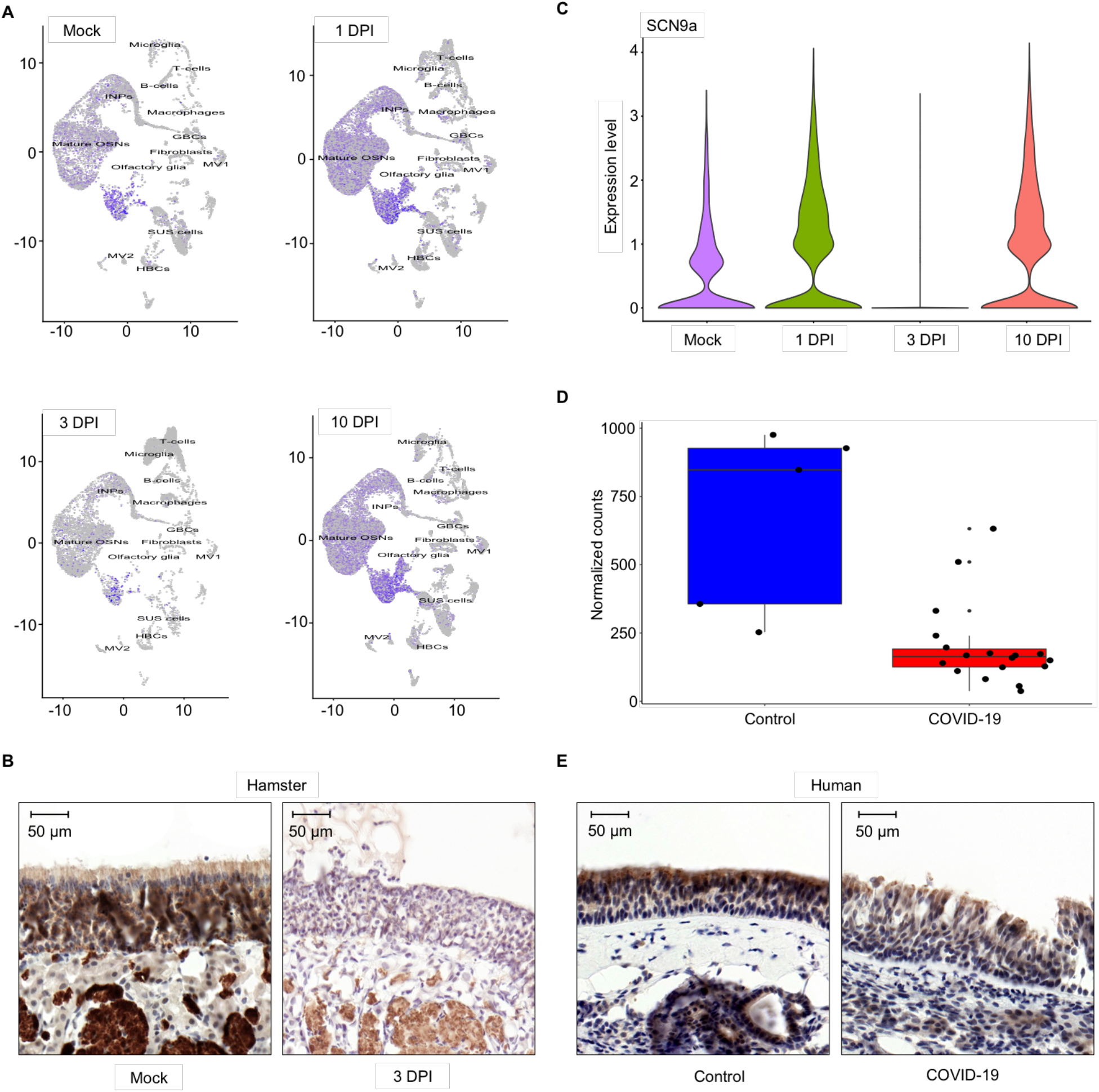
SCN9A gene expression in the olfactory epithelium of hamsters and humans infected with SARS-CoV-2. A) UMAP plots of SCN9A gene expression in different cell types of olfactory epithelium in mock and SARS-CoV-2 infected hamsters at 1, 3, 10 dpi. B) IHC slides of Na_V_1.7 expression in mock and infected hamster’s olfactory epithelium C) Violin plots of SCN9A gene expression in olfactory epithelium bulk tissues in mock and SARS-CoV-2 infected hamsters at 1, 3, 10 dpi. D) SCN9A gene expression in human OE tissues in control and SARS-CoV-2 infected cadavers. E) IHC slides of Na_V_1.7 expression in control and infected human olfactory epithelium

#### SARS-CoV-2 infection induces downregulation of the SCN9A gene in human OSN

We analyzed whether changes in SCN9A expression in ROEB observed in COVID-19 infected mice and hamsters is also observed in human OSNs post COVID-19 infection. The region of the cribriform plate located at the roof of the nasal cavity was resected, as a region with a high density of mature OSNs from 5 control and 18 infected human cadavers. We performed bulk RNA sequencing of olfactory epithelium tissues from the resected specimens. Cadavers were donated from patients of different sex (9 males, 16 females), with the median age of patients being 73 years (IQR 65-78), representing a variety of infection duration, hospital stay, treatment and postmortem interval. We have previously demonstrated that despite different postmortem times for collecting the samples, only minimal influence on the cellular constitution of tissues and immune cells in OE of humans could be observed [2]. SARS-CoV-2 was detected in all positive OE tissues with variations in the viral load [2]. SCN9A gene expression is 4-fold lower in SARS-CoV-2 infected OE tissues samples compared with healthy controls (p<0.001) (Figure 2d, e).

### Fluorescence Imaging of Na_V_1.7 expression in mouse ROEB using Tsp1a-IR800

Using widely available IVIS imaging system we could clearly visualize Na_V_1.7 expression in mouse ROEB using fluorescence imaging without need for any surgical intervention. The excitation was set to 750 nm and emission was set to 794 nm. We designed our probe to have these features because there is minimal background at these wavelengths to provide a highly specific signal. Because of superficial expression of Na_V_1.7 in the mouse ROEB, we were able to obtain images of the mice without need to expose the olfactory epithelium. Epifluorescence *in vivo* images in mice receiving IV injection with the imaging agent Tsp1a-IR800 generated high contrast between the ROEB and its surrounding regions. The radiant efficiency was significantly less in both mice injected with PBS and in mice injected with the unmodified peptide (blocking agent) in combination with the imaging agent (Fig. 3B). We observed a 150-fold increase in radiant efficiency compared to mice injected with PBS. To demonstrate specificity we co-administered our imaging probe with non-fluorophore labeled TSP1a and observed a 61-fold decrease in signal emanating from the mouse OE. For further confirmation, we resected the olfactory epithelium region of these mice and obtained regular tabletop fluorescent microscopy images of sectioned tissue (Fig. 4). The fluorescent microscopy images confirm the results of *in vivo* epifluorescence imaging and IHC. OSN and olfactory nerve bundles located in the lamina propria showed the most intense fluorescence. Normosmic mice and mice treated with blocking agent were negative for any signal (red), while only nuclear-associated blue staning was visible corresponding to nuclear staining by Hoechst dye.

**Figure 3.**
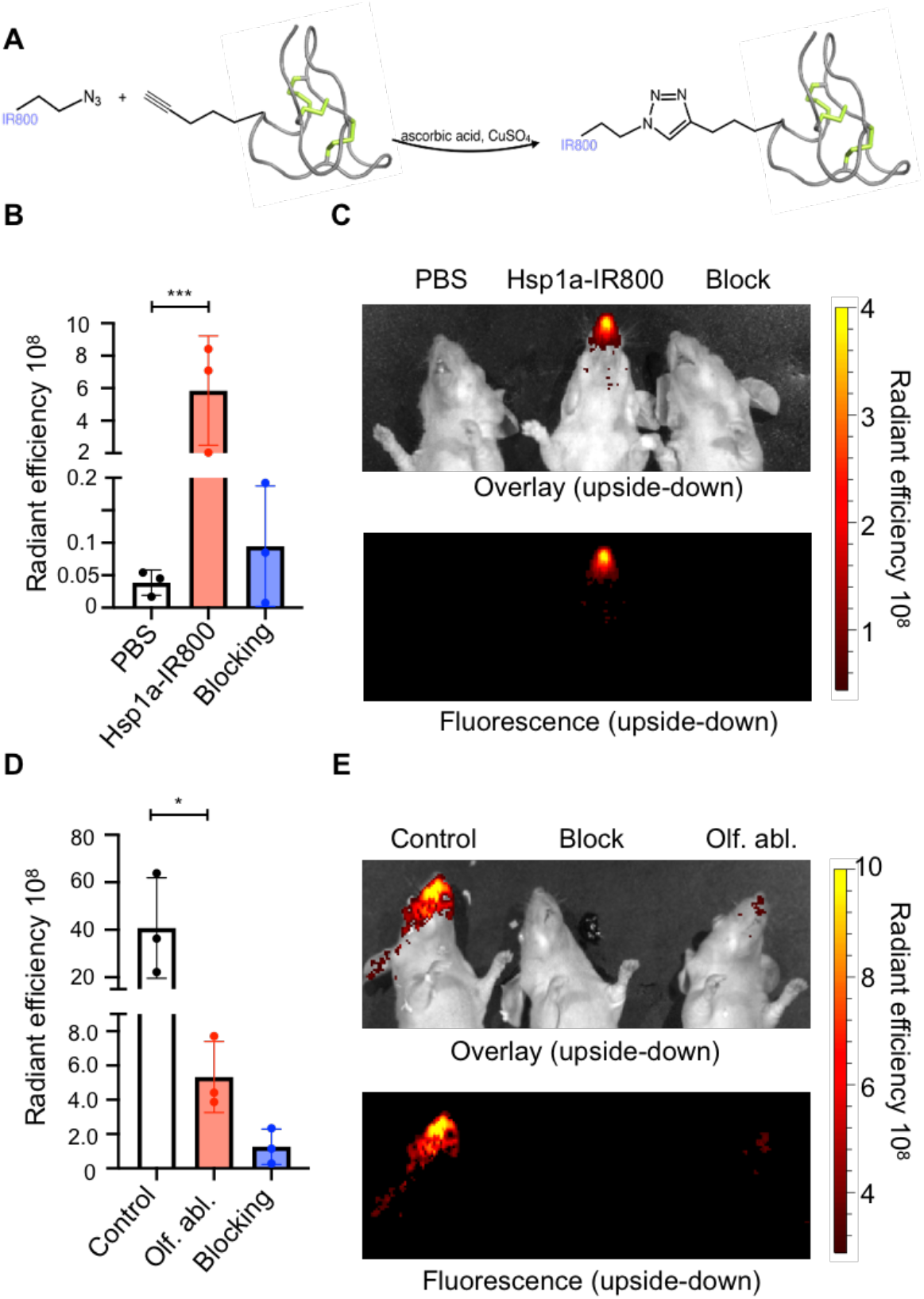
Tsp1a-IR800 accumulation in the ROEB in normosmic mice and mice with olfactory ablation. Chemical synthesis of Tsp1a-IR800. The IR800 fluorophore with an attached azido group reacts with an alkyne group on Tsp1a to yield the fluorescent imaging agent. (B-C) Epifluorescence images and fluorescent intensity quantification of animals injected with PBS, Tsp1a-IR800, and Tsp1a-IR800/Tsp1a blocking formulation, respectively. Images were taken 30 min after tail vein injection. (D-E) Epifluorescence images and fluorescence intensity quantification of normosmic control animals (injected with Tsp1a-IR800 or Tsp1a-IR800/Tsp1a blocking formulation) and mice with prior olfactory ablation with methimazole (injected with Tsp1a-IR800.) Images were taken 30 min after tail vein injection. (* p ≤ 0.05; ** p ≤ 0.01; *** p ≤ 0.001; **** P ≤ 0.0001).

**Figure 4.**
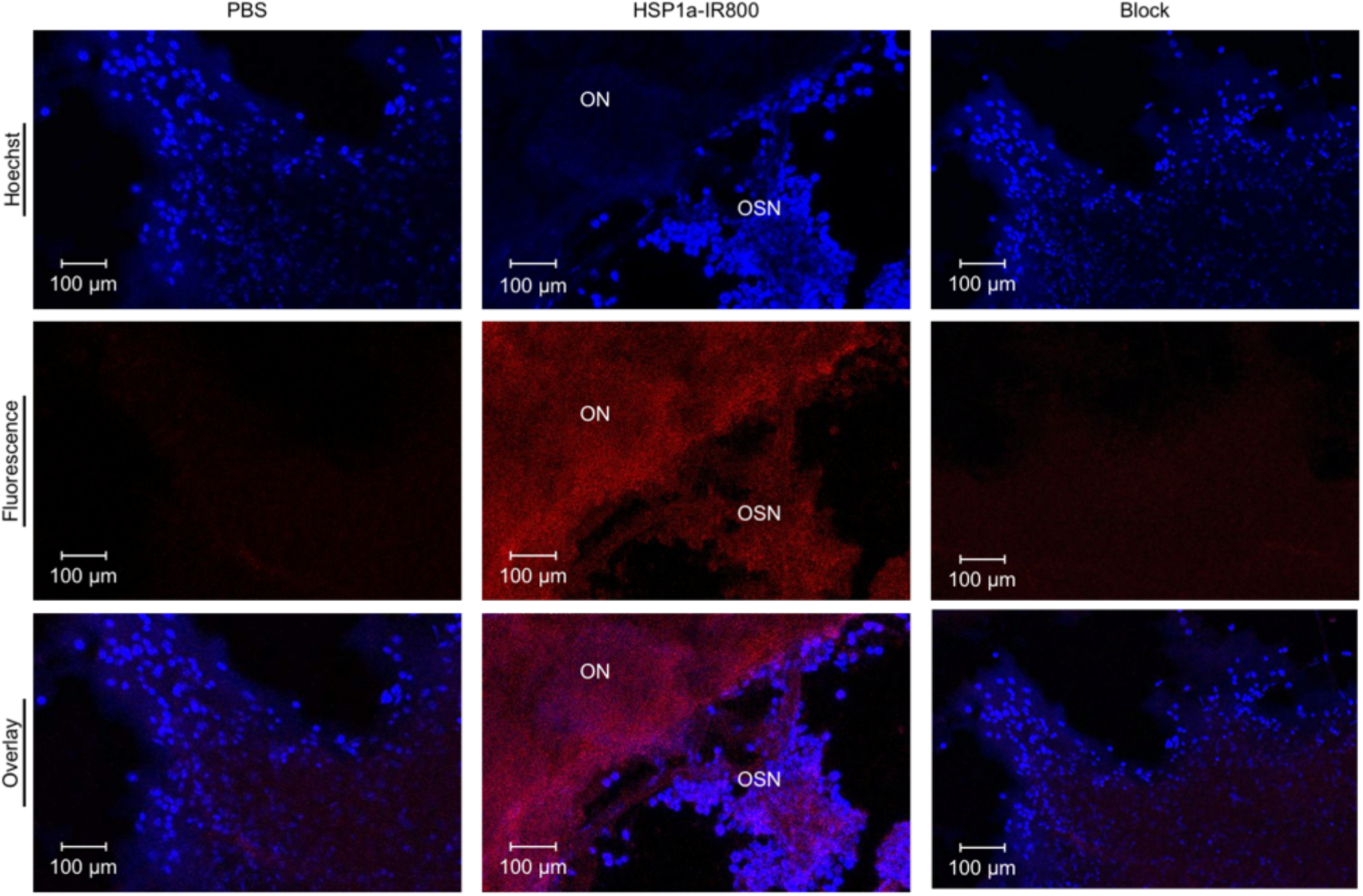
Fluorescent confocal microscopy images of the olfactory epithelium of animals injected with PBS, Tsp1a-IR800, and Tsp1a-IR800/Tsp1a blocking formulation, respectively. (ON – olfactory nerve (bundles), OSN – olfactory sensory neurons.)

### Chemically induced Anosmia can be monitored using fluorescence imaging with Tsp1a-IR800

Mice treated with methimazole that is known to cause damage to olfactory nerves had visibly diminished fluorescence signal (radiant efficiency) compared to normosmic control mice when imaged with Tsp1a-IR800. The ROEB of methimazole-treated mice had a 8-fold decrease in radiant efficiency compared to normosmic mice (Fig. 3D&E). The average radiant efficiency of the olfactory region of normosmic mice imaged from both sides were 4.08E+09 (s.d. 2.11E+09), compared to 5.33E+08 (s.d. 2.08E+08) for mice with an olfactory ablation (unpaired *t*-test, p=0.045). *Ex vivo* images showed a similar statistically significant difference between mouse groups in the ROEB (do we have supplemental data figure?). The heart and kidneys are the only internal organs with higher fluorescence signals compared to normosmic mice and mice treated with blocking agent.

### Time to completion of the buried food test correlates with the radiant efficiency of the ROEB in normosmic and olfactory-ablated mice

Mice with olfactory ablation needed a significantly longer time than normosmic mice to find buried food (Fig. 5B). Normosmic mice were able to find buried food in less than 30 s, compared with average time of 135 s for olfactory-ablated mice (p<0.001). Furthermore, there was an inverse correlation between the <u>Tsp1a-IR800</u> radiant efficiency and the time required to find buried food (Pearson correlation coefficient r = – 0.79, n=10, p=0.0056; Fig. 5C).

**Figure 5.**
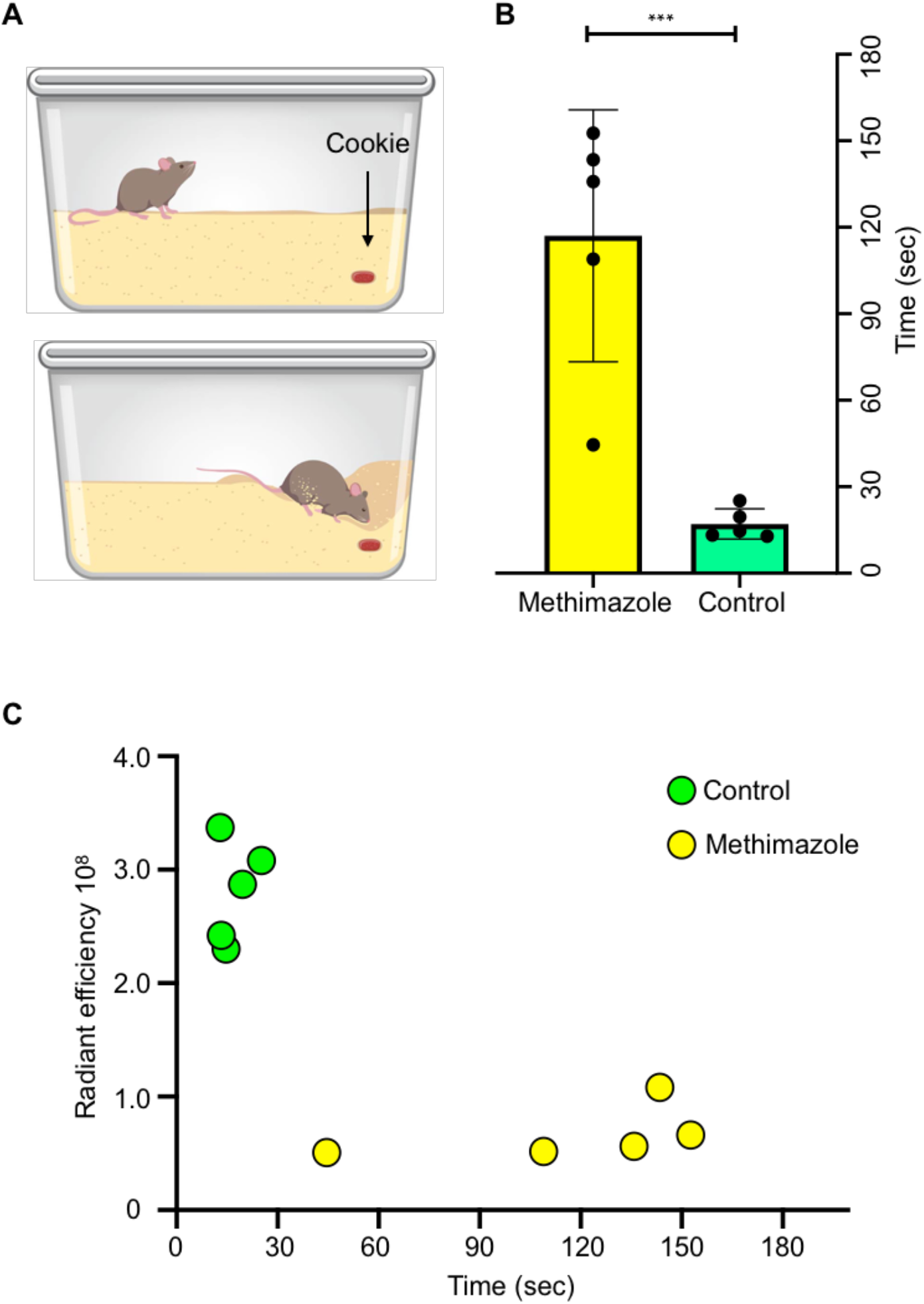
Buried food test. (A) Schematic illustrating mouse cage with a cookie buried in the upper right corner of the cage. (B) Bars show time (seconds) spent for mice treated with PBS (n=5) or methimazole (n=5) to find the buried food. (C) Correlation of <u>Tsp1a-IR800</u> radiant efficiency at ROEB and time in buried food test. (* P ≤ 0.05; ** P ≤ 0.01; *** P ≤ 0.001; **** P ≤ 0.0001.)

#### Na_V_1.7 expression in olfactory epithelium of non-human primates (NHP) can be detected using fluorescence imaging of Tsp1a-IR800 with clinically approved near-infrared (NIR) endoscopy systems

To validate the clinical potential of our approach of the imaging Na_V_1.7 expression as a surrogate marker for sense of smell, we performed experiments in non-human primates (NHP) using the Quest NIR imaging system that is approved for clinical use. After IV administration of Tsp1a-IR800, tissues were harvested. The bright fluorescence is clearly visible over the olfactory epithelium, and weak or no fluorescence is detected over the muscle, olfactory bulb, and brain. The fluorescence intensity of olfactory epithelium was significantly higher compared to all other measured tissues (Fig. 6A&B). Furthermore, we performed fluorescence microscopy of the dissected non-human primate tissues (Fig. 6C). The fluorescence signal from the NHP olfactory epithelium was clearly brighter compared to other tissues. It is important to realize that the olfactory bulb expresses Na_V_1.7 but the blood brain barrier (BBB) prevents the entry of our imaging agent in the current setting. For the current application, this is a significant advantage because it reduces potential background signal from Olfactory bulb.

**Figure 6.**
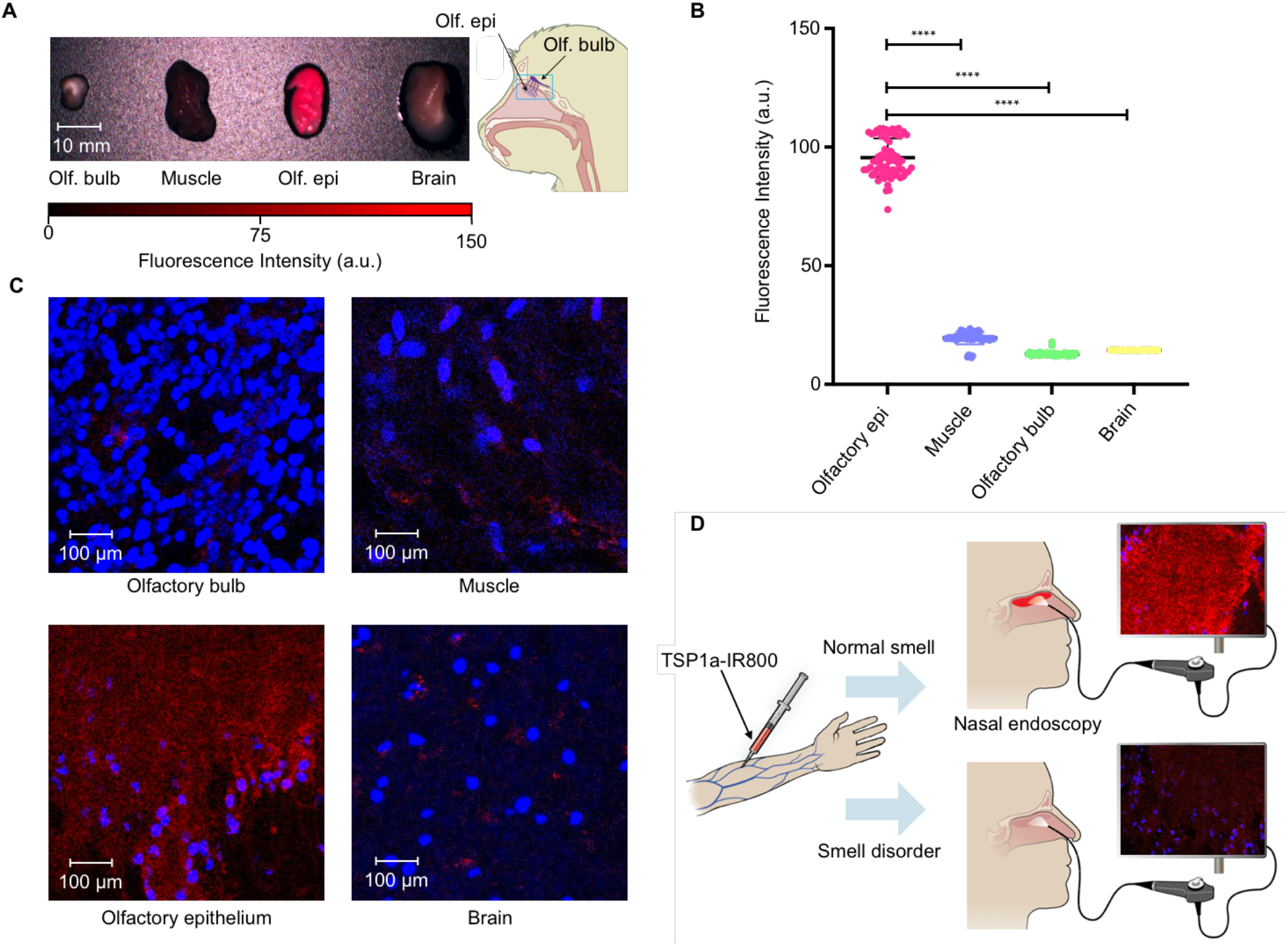
Imaging of olfactory epithelium in non-human primates. (A) Images were taken using Quest system (approved for clinical use) of the olfactory bulb, muscle, olfactory epithelium, and the brain of the NHP after intravenous injection of TSP1a-IR800 (B) Quantification of the near infra-red fluorescence intensity using authorized Quest software. (C) Fluorescent confocal microscopy images of the olfactory bulb, muscle, olfactory epithelium and brain tissues of the same NHPs (D) Schematic depiction of the potential use of the TSP1a-IR800 in the physician’s office setting using the Quest or other vendor NIR fluorescence imaging systems. (* P ≤ 0.05; ** P ≤ 0.01; *** P ≤ 0.001; **** P ≤ 0.0001.)

## Discussion

We describe development of a novel semi-quantitative diagnostic method based on selective targeting of Na_V_1.7 expression in olfactory epithelium for detection of smell disorders induced by different insults and inflammatory conditions such as COVID-19. The fluorescence signal emanating from mouse OE shows decreased intensity when mouse have Anosmia. There was inverse linear relationship between signal intensity and the degree of damage. Therefore we believe it has potential to serve as an objective guide to assess disease progression as well as treatment response both in animal and human subjects. The ability to asses olfaction in animal models opens new avenues for drug development and testing. All current diagnostic methods are based on a smell perception of the afflicted patients and are subjective by design. Despite being standardized and validated, these methods are culturally specific and individually subjective [19]. Certain odors may not be familiar to those outside the geographic region where the test had been developed. Moreover, odor identification methods only can diagnose the presence of a problem, but not the site of the damage or etiology. The only existing method that can do both is an invasive surgical biopsy that can lead to tissue destruction. However, the invasive nature of the procedure limits its use in multiple ways. Furthermore, in animal studies, only behavioral experiments are available to measure smell loss, which cannot be repeated often, can give non consistent results and will not detect early changes. Because of the objective nature of our approach, we believe it opens a completely new paradigm to detect olfaction in animal subjects.

A new diagnostic method, that is non-invasive, can be easily and repeatedly performed during a routine exam, and can help to diagnose the level of damage is crucially important. The absence of a good diagnostic method hampers any efforts in developing a novel treatment strategy that is tailored to a specific cause. You cannot treat central and peripheral causes of anosmia the same way. Moreover, the method should be able to detect the restoration process before noticeable improvement in sense of smell [4].

We have identified that smell loss due to chronic inflammation in the nasal cavity or viral infection all are accompanied with the diminished expression of Na_V_1.7 sodium channels. We observed loss of Na_V_1.7 channel expression following olfactory ablation using methimazole (Fig. 1). The expression of Na_V_1.7 channels was substantially reduced in the OSN, and nerve bundles located in lamina propria, whereas it was less affected at the level of olfactory bulbs.

Further, we assessed whether COVID-19-related olfaction loss is also accompanied with the loss of Na_V_1.7 sodium channels. COVID-19-related smell disorders are possibly caused by multiple mechanisms. There is a direct tropism of the virus to sustentacular and microvillar cells covering the OSN, inflammatory damage to OSN, as they are exposed to environmental factors, and inflammation involving focal mucosal swelling and obstruction to airways [20]. Recently, our collaborators and co-authors on this publication described that downregulation of odor detection pathways as a potential cause of COVID-19 induced anosmia [2]. We found out that together these mechanisms cause diminished Na_V_1.7 channel expression in the OSN, and olfactory nerve bundles located in the nasal cavity as shown by IHC and RNA sequencing on tissues from mice, hamsters, and humans.

Our compound Tsp1a-IR800 selectively binds to Na_V_1.7 sodium channel, making it ideal candidate to measure smell perception using fluorescent measurement technics. We tested our imaging agent using a mouse model in which olfactory ablation was achieved via methimazole injections. This damage follows a similar sensorineural smell loss pattern seen in upper-respiratory viral infection or chronic inflammation due to seasonal allergies. After iv injection of the compound, we observed diminished radiance efficiency in mice with ablated OE, compared with the normosmic mice. Further, we have detected that the extent of Na_V_1.7 expression as measured by radiant efficiency of the Na_V_1.7 imaging agent was inversely correlated with the time mouse spent finding food in a buried food test.

The difference between small mammals and humans can be substantial, and to validate the possibility to image olfaction in humans, we injected the compound to a healthy non-human primate. For imaging of fluorescence, we have used clinically approved NIR imaging machine. We observed bright fluorescence from the olfactory epithelium, and the fluorescence measurements were significantly higher compared with surrounding tissues.

As described above, it is crucial to develop a non-invasive, fast, and objective method to diagnose smell disorders. Intravenously injected Tsp1a-IR800 selectively accumulates in OSN and olfactory nerve bundles located in the nasal cavity. Existing flexible endoscopes from several medical technology companies (Quest, Stryker, Olympus etc.) can detect the IR wavelength emitted by Tsp1a-IR800 (Fig. 5D). Due to the minimally invasive nature of the nasal cavity scope, the correlation of fluorescence with the magnitude of smell loss the method can be used as a treatment monitoring tool, and in the development of therapeutic treatments. In addition, the method can be widely used in an experimental setting, to objectively measure smell in animals replacing the need to perform behavioral experiments and streamline the development of new therapeutics or investigating diseases linked with smell disorders.

## Declarations

### Author Contributions

D.A., J.G., T.R., S.P. and N.P. conceived the study and designed the experiments. D.A. J.G, T.V., P.D.S.F, S.R., S.J, A.O., L.C. and T.R. carried out the experiments and collected the data. J.G., C.Y.C., G.K. and T.R. produced Tps1a-IR800. D.A., N.P. and T.R. analyzed the data, D.A., J.G., N.P. and T.R. conducted statistical analysis of the data. D.A., J.G., N.P. and T.R. primarily wrote and edited the manuscript. All the authors reviewed and approved the manuscript.

## Acknowledgements

The authors thank the support of Memorial Sloan Kettering Cancer Center’s Animal Imaging Core Facility and Molecular Cytology Core Facility. We especially thank Eric Chan for the help with the Image J plugins for Na_V_1.7 quantifications. We also thank Terry Helms for creating illustrations.

